# Evolutionary dynamics and diversification in changing environments

**DOI:** 10.1101/590406

**Authors:** Evgeniia Alekseeva, Michael Doebeli, Iaroslav Ispolatov

## Abstract

We use adaptive dynamics models to study how changes in the abiotic environment affect patterns of evolutionary dynamics and diversity in evolving communities of organisms with complex phenotypes. The models are based on the logistic competition model and environmental changes are implemented as a temporal change of the carrying capacity as a function of phenotype. In general we observe that environmental changes cause a reduction in the number of species, in total population size, and in phenotypic diversity. The rate of environmental change is crucial for determining whether a community survives or undergoes extinction. Until some critical rate of environmental changes, species are able to follow evolutionarily the shifting phenotypic optimum of the carrying capacity, and many communities adapt to the changing conditions and converge to new stationary states. When environmental changes stop, such communities gradually restore their initial phenotypic diversity.

## Introduction

Over the past decades, the issue of the impact of changing environmental conditions on species and ecosystems has gained increasing prominence, particularly in the context of global warming (Griggs and Noguer 2002). Recent estimates have shown that at the current rate of global warming, one of six species will become extinct (Urban 2015), and empirical evidence supports this finding (Maclean and Wilson 2011). Already there are species whose extinction occurred as a result of climate change. For example, the sea level rise has destroyed the habitat of mosaic-tailed rat *Melomys rubicola* and individuals of this species have not been seen since 2009 (Gynther et al. 2016). Many species, such as polar bears (*Ursus maritimus*), experience ecological stress. For hunting, this species relies on sea ice, where seals, their primary source of food, rest and breed. Reduction of ice surfaces forces polar bears to overcome long distances by swimming and thus strongly affects the balance between anabolism and catabolism. Several studies detected muscle atrophy and weight loss in polar bears because of starvation (Obbard et al. 2016a,b; Whiteman et al. 2017; Griffen 2018; Pagano et al. 2018).

However, the majority of ecosystems are characterized by extensive adaptability. While changing environmental factors often lead to diversity reduction, in general many ecosystems will likely survive. For example, this is currently observed in coral reefs. Increasing temperature and acidification of the ocean water affect the symbiotic relationships between corals and microalgae in such a way that corals expel their endosymbionts and bleach (Pogoreutz et al. 2018). Without benefits of symbiosis, corals experience higher mortality, become more sensitive to diseases, and decline. However, because strains of both host and endosymbiont vary in their sensitivity to higher temperatures, some of them can form a thermo-tolerant symbiosis (LaJeunesse 2002; Baker 2003; Little et al. 2004; Smith et al. 2017; Grottoli et al. 2018). Such examples already exist in zones **with extreme temperatures, such as the Arabian / Persian Gulf (PAG)** (Baker et al. 2004), hence this symbiotic community can adapt, in principle, to changing environments despite a decrease in diversity of its participants.

The challenges imposed by changing environments vary widely between ecosystems and between species within ecosystems, as e.g. the warming climate induces a diverse spectrum of interconnected changes in environmental conditions and weather patterns. Besides global climatic changes, there are numerous other examples how anthropogenic activity disturbs ecosystems locally by environmental pollution, poaching, modification of geographical landscapes and many others factors (Laskar et al. 2016; Scheffers et al. 2019). Adaptation to changing environments is also an important topic of research in the context of preventing the development of antibiotic and drug resistance. So, in one way or another, biological populations frequently face changing environment, which is an important force in their evolution.

Adaptation to environmental changes has been the subject of both experimental and theoretical research. A nice example of an experimental study of evolution in artificially created changing environment is the work on gradual bacterial adaptation to increasing doses of antibiotic on a giant Petri dish (Baym et al. 2016). Another example of experimental adaptation to a changing environment was observed in a study of phytoplankton biodiversity in increasingly warm water (Yvon-Durocher et al. 2015). However, the slowness of evolutionary processes on human timescales sets strong restrictions of what can be done in such experiments. Free of this limitation, theoretical studies are by far more numerous. For example in (Botero et al. 2015), the authors found that certain types of climatic changes force populations to cross “tipping points” and to switch from one adaptive strategy to a qualitatively different one, which often leads to extinction despite the successful adaptation in the context of the previously used strategy. Another theoretical study has revealed the influence of genetic variance and spatial dispersal on the success of a given number of competing species subjected to changing conditions (Norberg et al. 2012). The combination of high genetic variance and low spatial dispersal is the most conducive to adaptation and survival of the species under the effect of climatic change. In (Northfield and Ives 2013), the authors have investigated how coevolution in pairs of species with various types of ecological interactions affects the process of adaptation to environmental changes. They argued that types of coevolution with conflicting interests help species to adapt, whereas types of coevolution with non-conflicting interests enhance the detrimental effect of climatic changes.

In this work we take a somewhat different look at the influence of environmental changes on an evolving system. As is widely reported, the problem with environmental changes is often not so much the actual state of the environmental variable, such as the global temperature or the *CO*_2_ concentration, but the high and previously unseen rates at which these variables change. Thus, in this work we investigate how an ecosystem, modelled as a community of interacting and evolving species, reacts to environmental changes of various rates. We focus on the particular case of competing species, ignoring for now other ecological interactions, and consider a diversifying community described by a logistic competition model in which the competition between individuals is controlled by several phenotypic traits. Previously it was shown that in such systems, the dimension of phenotype space affects diversification, with higher dimensions leading to higher diversity (Doebeli and Ispolatov 2017). As a community diversifies from low numbers of species, the rates of evolution and of diversification slow down as the community reaches a saturation level of diversity that the environment can sustain. Higher rates of evolution and diversification can be reactivated only when the level of saturation decreases, which can happen with aromorphosis and an extension of the phenotypic space into higher dimensions, or through catastrophic events, leading to mass extinction (Ispolatov, Alekseeva, et al. 2019).

However, it is not known which ecological and evolutionary processes unravel in such a system when external intervention, such as ongoing climate changes, persists indefinitely. A naive qualitative guess (which turns out to be correct) would be that if the rate of change associated with such an intervention is much smaller than some intrinsic adaptation rate of all species, the relative phenotypic distribution of the species would remain almost intact and all species would synchronously follow the environmental change. Yet it is hard to predict even qualitatively what happens when the rate of environmental changes increases, apart from the ultimate extinction of all species when environmental change is very fast. We therefore performed a systematic study of various ecological and evolutionary indicators of the evolving communities for a wide range of rates of environmental changes.

## Methods

### The Model

Following (Doebeli and Ispolatov 2017; Ispolatov, Alekseeva, et al. 2019), we study a system that can be populated by a varying number of phenotypic species, each defined by its phenotype **x** = (*x*_1_, …, *x*_*D*_) in *D*-dimensional phenotype space. In monomorphic communities consisting of a single species with phenotype **x**, the population size of that species at ecological equilibrium is given by the carrying capacity function *K*(**x**), which is assumed to have the following form:

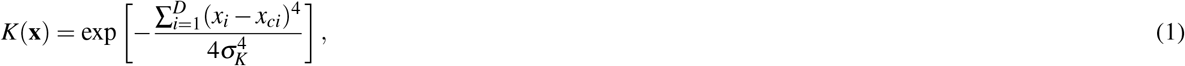

where *σ*_*K*_ determines the width of the carrying capacity. This function has its maximum at the point that we call the centre of the carrying capacity (CCC) **x**_*c*_ = (*x*_*c*1_, …, *x*_*cD*_). Thus, the population size of a monomorphic population is maximal if that population’s phenotype is equal to **x**_*c*_.

Competition between two phenotypes **x** and **y** is described by the competition kernel *α*(**x**, **y**), so that the competitive effect of **x** on **y** is given by

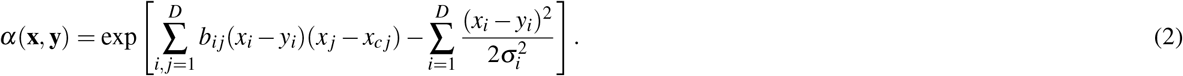

There are two terms in the exponent of the competition kernel. The first one represents the simplest non-symmetric contribution to the competition that may result in complex evolutionary dynamics (Doebeli and Ispolatov 2014; Ispolatov, Madhok, et al. 2016; Doebeli and Ispolatov 2017). Since we expect that the evolutionary dynamics would unravel around the CCC (positioned at zero in (Doebeli and Ispolatov 2014; Ispolatov, Madhok, et al. 2016; Doebeli and Ispolatov 2017)), we explicitly introduce the coordinates of CCC **x**_**c**_, replacing *x*_*j*_ used in (Doebeli and Ispolatov 2014; Ispolatov, Madhok, et al. 2016; Doebeli and Ispolatov 2017) by (*x*_*j*_ −*x*_*cj*_). The second term in the exponent is the usual Gaussian competition kernel with width *σ*_*i*_, reflecting the fact that species that are closer phenotypically compete more strongly with each other than species that are farther apart in phenotype space. In our simulations we used *σ*_*K*_ = 1 and *σ*_*i*_ = 1/2 to ensure that the system is able to diversify from the initial state of one species to a community of coexisting phenotypes (Ispolatov, Madhok, et al. 2016; Doebeli and Ispolatov 2017). Also as in (Doebeli and Ispolatov 2017), the coefficients *b*_*ij*_ of the non-symmetric part of the competition kernel were chosen randomly from a Gaussian distribution with width 1 and zero mean (see the end of this section for how this was implemented to obtain the simulation results).

Assuming that a community comprises *m* species with phenotypes **x**_*r*_ = (*x*_*r*1_, …, *x*_*rD*_) in *D*-dimensional phenotype space, where *r* = 1, …, *m* is the species index, the ecological dynamics of the density *N*_*r*_ of species *r* is given by the logistic equation

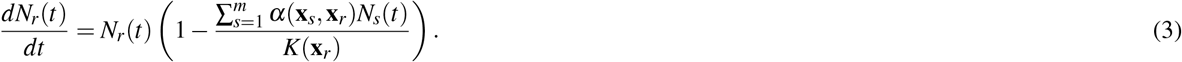

In the framework of adaptive dynamics (Geritz et al. 1998; Diekmann 2002), evolution occurs when species generate rare mutants with phenotypes that are close to but distinct from the parent phenotype. Mutants compete with the resident population for resources and try to invade it with a per capita growth rate given by the invasion fitness function 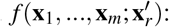:

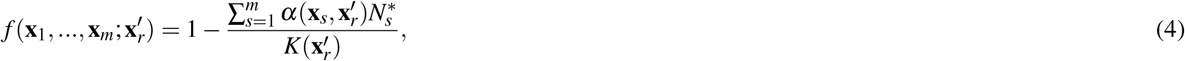

where 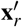 is the phenotype of a mutant occurring in species *r*. By differentiating the invasion fitness with respect to the mutant phenotype and evaluating at the resident phenotype, one obtains the selection gradient *S*_*r*_ with components *S*_*ri*_:

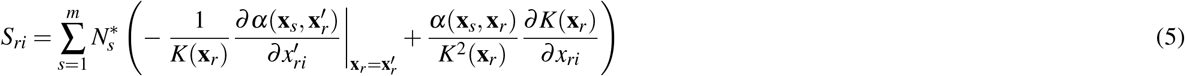

Adaptive dynamics of the phenotypes of a *m* species in a community is determined by the selection gradient and by the mutational variance-covariance matrix, which describes the rate and size of mutations occurring in each species. For simplicity, we assume here that this matrix is diagonal, and that the elements corresponding to each species are proportional to the current population size of that species. The adaptive dynamics of each phenotypic component *x*_*ri*_, *i* = 1, …, *D* and *r* = 1, … , *m* is then given by

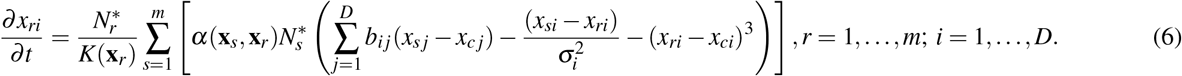

Further details of the model, including the procedure allowing species to diversify, are presented in the Appendix, Section 1. So far, this model has been defined in the same way as the one in (Ispolatov, Madhok, et al. 2016; Doebeli and Ispolatov 2017; Ispolatov, Alekseeva, et al. 2019). Here, however, we introduce environmental change by assuming that over time, new phenotypes become optimal for the current state of the changing environment. Since the optimal phenotype in our model is defined by the position of the CCC in the phenotypic space, the environmental changes can be implemented as the motion of the CCC and thus the carrying capacity itself in the phenotypic space, **x**_*c*_ = **x**_*c*_(*t*) = **V**_*C*_*t*. Here **V**_*C*_ is a vector of a given length *V*_*C*_, which defines the absolute rate of environmental change, as well as its direction, which is the direction in phenotype space along which the part of selection gradient generated by the carrying capacity is strongest. While the maximum of the carrying capacity function moves in phenotype space at a constant rate *V*_*C*_, the general shape of the carrying capacity function, and in particular its width *σ*_*K*_, are assumed to stay the same in the moving frame of reference in phenotype space.

In the simulations, environmental change starts once an evolving community has reached a stationary state in the evolutionary dynamics with constant environment (Fig. S1), as described in (Doebeli and Ispolatov 2017; Ispolatov, Alekseeva, et al. 2019). In the following we denote this time as *t*\*. In our simulation we prepare 30 replicate simulations for every phenotypic dimensionality *D* = 1, 2, and 3, each having a distinct randomly chosen set of coefficients *b_i j_* and initial conditions. Each replicate is evolved for time *t*\* to its steady state following the procedure outlined in Appendix, Fig. S1. Then, in every dimensionality *D* = 1, 2, and 3 and for each magnitude of *V*_*C*_, we perform 30 simulation runs, one for each steady state replica. In each run, a new random direction of **V**_**C**_ is chosen for the same magnitude *V*_*C*_. Once started, the environmental change occurs at a constant rate *V*_*C*_ for *t*\* time units. The results for each *D* and *V*_*C*_ are averaged over those 30 runs, producing statistical data shown in Fig. 2 for the final state and in Fig. 3 for the time course of adaptation (Appendix, Section 1).

**Figure 1.**
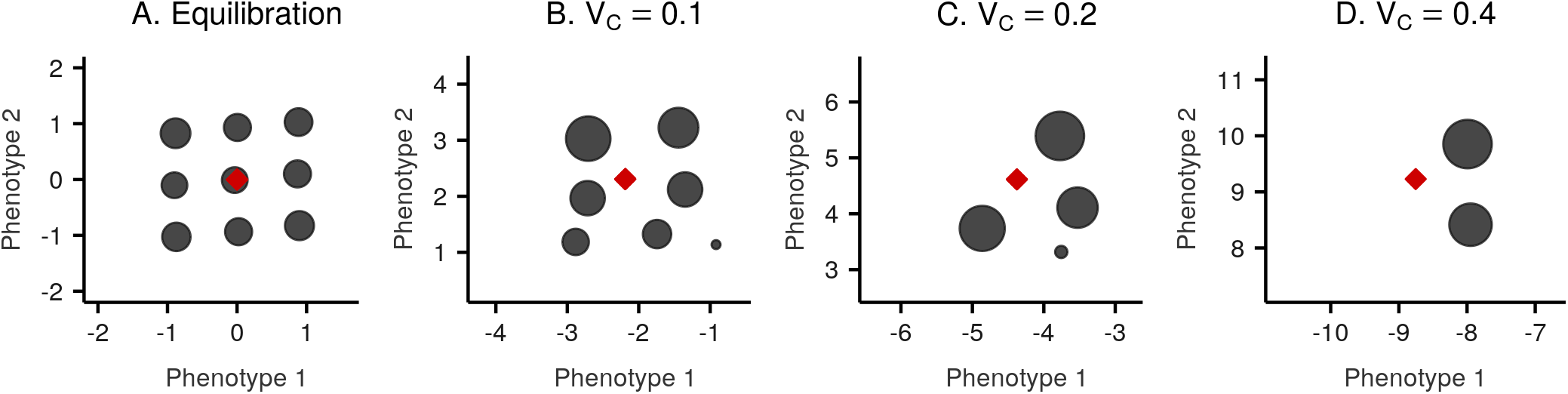
Snapshot of the saturated diversification (**A**) and adaptation to environmental changes of various rates (**B**, **C**, **D**) in two-dimensional phenotypic space, *σ*_*K*_ = 1, *σ*_*i*=1,…,*D*_ = 0.5; processes that led to these configurations can be seen in corresponding videos <u>here</u>. Dark grey circles show the location of different species in phenotype space, with the size of the circles representing the populations size, the red rhombus shows the location of the CCC. The coefficients *b*_*ij*_ and the initial conditions can be found in Appendix, section 2.

**Figure 2.**
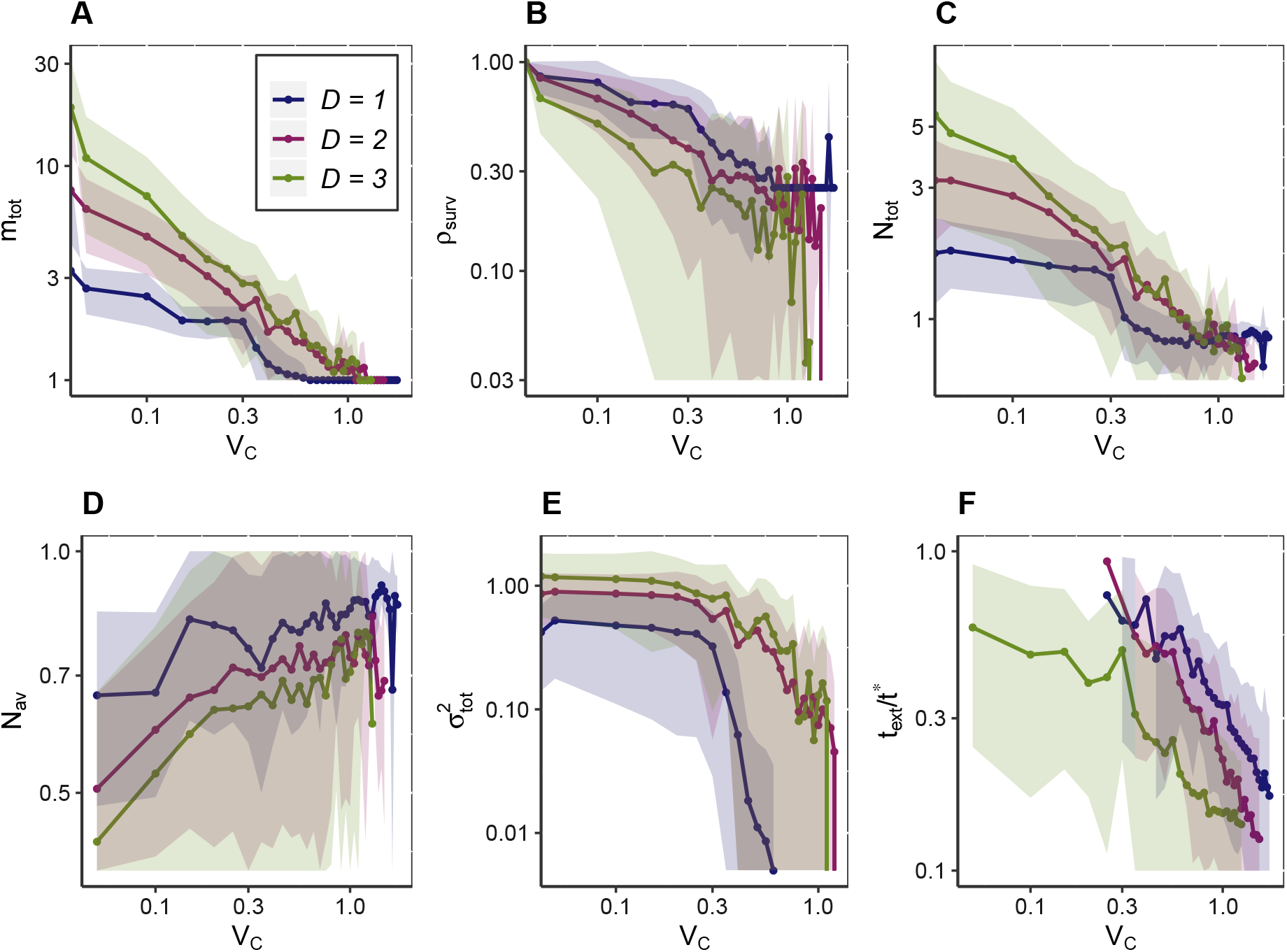
The total number of species *m*_*tot*_ (**A**), the fraction of species that survived *ρ*_*surv*_ (**B**), the total community population *N*_*tot*_ (**C**), the average per species population *N*_*av*_ (**D**), the phenotypic variation across the community 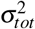 (**E**), and the normalized time to total extinction *t*_*ext*_/*t*\* (**F**) vs. the speed of environmental change *V*_*C*_. For each value of *V*_*C*_ all these quantities with the exception of the extinction time were averaged over 30 systems, after each of them evolved under the changing environment for time *t*\* and reached new quasi-stationary states. The extinction time was measured for communities in which all species went extinct before *t*\*. Colours indicate the dimensions of phenotype space: blue for *D* = 1, red for *D* = 2, and green for *D* = 3. Values of *t*\* for each dimensionality of the phenotype space are presented in Fig. S2 in the Appendix. To reveal the power-law-like nature of many dependencies, all figures are presented in log-log scale with shadows around lines indicating standard deviations.

**Figure 3.**
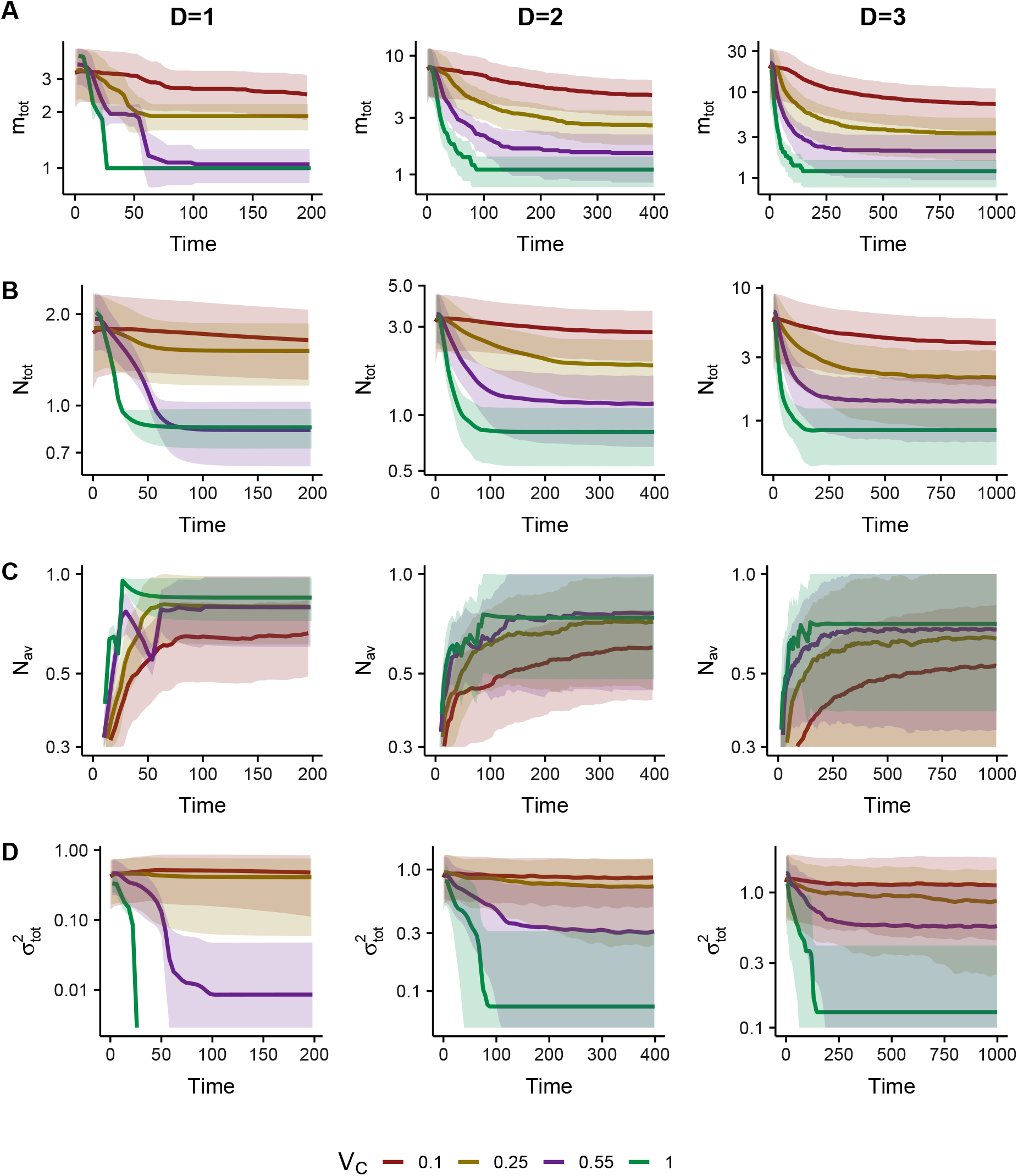
Dynamics of **A**: average number of species *m*_*tot*_ ; **B**: total population *N*_*tot*_ ; **C**: average population of a species *N*_*av*_ and **D**: phenotypic variation across the community, 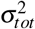 as a function of time after the onset of environmental change for different values of *V*_*C*_. Figures are presented in semi-logarithmic scale.

### Measuring properties of the system

To analyze the evolving communities, we measured the following system properties at *t*\*:

1. number of species in the system *m*_*tot*_.
2. total population of the system 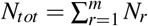.
3. average population of a species 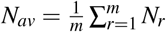.
4. phenotypic diversity of the system 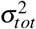 as the average square distance of the phenotypic coordinates of all clusters weighted by their population sizes around the centre of mass of the system 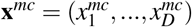, where

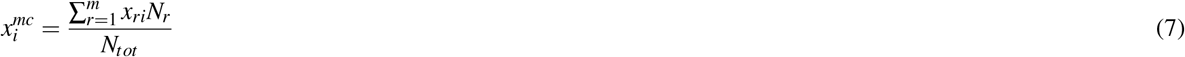

and

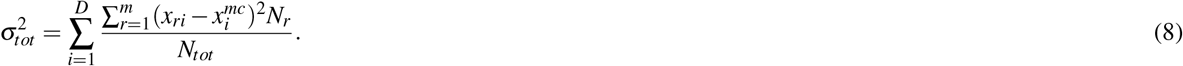 The quantity 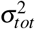 reflects how widely the phenotypes of the various species are separated from the centre of mass of the system. It is a measure of the phenotypic diversity in a community, but should not be confused with the number of species, since the phenotype distribution in systems with smaller numbers of species can nevertheless have a higher variance if the fewer species are more spread out in phenotype space.
5. fraction of survived species

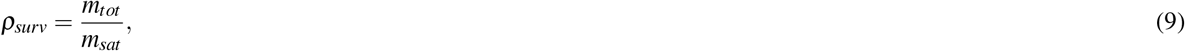

where *m*_*sat*_ is the number of species in saturated communities before the initiation of environmental change.
6. time to extinction *t*_*ext*_; extinction occurs when all species of the system go extinct before time *t*\* after the onset of environmental change.

There are two basic ways in which the above quantities can be calculated to illustrate system behaviour. First, they can be evaluated at the final time *t*\* after the onset of environmental change for many different systems with the same control parameters (e.g., the same *V*_*C*_). For example, one can calculate the average number of coexisting species at time *t*\*, i.e., the average number of coexisting species at the evolutionary quasi-stationary state, by averaging *m*_*tot*_ at *t*\* for many different systems. Second, these quantities can be studied as a function of time in any given simulation run, usually starting from the initiation of the movement of CCC, i.e., from the beginning of the environmental change. For example, before starting the movement of CCC, *m*_*tot*_ = *m*_*sat*_, i.e., the number of species at saturation. Once CCC starts to move, *m*_*tot*_ may change, depending on the effect of environmental change on a previously saturated community.

## Results

We analysed the adaptation of 30 saturated systems, properties of which were described previously (Doebeli and Ispolatov 2017, Ispolatov, Alekseeva, et al. 2019). For all those systems, the environmental change in the form of a moving CCC either forces the system to adapt and converge to a new quasi-stationary state, or it leads to extinction of the whole community.

### Adaptation to the CCC motion and convergence to a new quasi-stationary state

When the rate of environmental change is not too fast, after a transitory adaptation the system usually converges to a new quasi-stationary state where it follows the changing environment. Properties of such a quasi-stationary state vary depending on the rate of environmental changes. In Fig. 1 there is a two-dimensional visualisation of newly formed stationary configurations of the same community, adapting to different rates of *V*_*C*_.

For each value of *V*_*C*_, the final number of species *m*_*tot*_ , the fraction of surviving species *ρ*_*surv*_ and the total population size *N*_*tot*_ (Fig. 2 A:C) always decrease with increasing speed of environmental changes, *V*_*C*_. For a given value of *V*_*C*_, the number of species *m*_*tot*_ and total population *N*_*tot*_ begin to decrease shortly after the onset of environmental change and after some time stabilize (despite the ongoing CCC movement), as illustrated in Fig. 3 A and B. The higher the *V*_*C*_ is, the faster this decrease occurs. Conversely, the average per species population *N*_*av*_ in quasi-stationary stage increases with increasing *V*_*C*_: as more species go extinct during the process of adaptation, the surviving ones become more ecologically successful due to experiencing less competition (Fig. 2D and Fig. 3C).

The average value of phenotypic diversity, 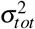, decreases for larger *V*_*C*_ as well (Fig. 2D), since larger rates of environmental change cause the loss of more species. Over the course of an individual simulation run, after the onset of environmental change 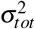 first decreases with time and then approaches a stationary state. For very small values of *V*_*C*_, phenotypic dispersion may stay at the level of saturated system without environmental change or even exceed it despite a reduction in the total number of species *m*_*tot*_ (Fig. 3D). Such behaviour of 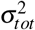 indicates that phenotypic spread of the surviving species around the centre of mass becomes wider.

Typically, the time required to reach a new evolutionarily stationary state is small relative to the saturation time *t*\*. The higher the rate of environmental change *V*_*C*_ is, the less time is required. The timescale of Fig. 3 is two times shorter than the timescale required for initial equilibration (Appendix, Fig. S2).

Even for small values of *V*_*C*_, the entire evolving community can go extinct. Such events become more likely for larger rates of environmental changes *V*_*C*_. Our simulations indicate that for any fixed set of parameters, there is a value of *V*_*C*_ for which the extinction becomes certain. We call this value the extinction threshold. The fraction of surviving communities as a function of *V*_*C*_ and the extinction threshold value of *V*_*C*_ are shown in Fig. S3 of the Appendix. Generally, the higher the rate of environmental change, the more likely extinction occurs (Fig. S3C). Times to extinction *t*_*ext*_/*t*\* become shorter for larger values of *V*_*C*_ (Fig. 2F).

### The influence of phenotypic complexity on adaptation and extinction

Having more phenotypic dimensions complicates the adaptation to environmental changes. The surviving species in low-dimensional phenotype space have larger populations *N*_*av*_ (Fig. 2D) and even the overall population of the community *N*_*tot*_ of a low-dimensional system is larger than that of a higher-dimensional one for high rates of *V*_*C*_. Furthermore, the time required for the community to reach the quasi-stationary state becomes larger for larger phenotypic complexity (Fig. 3).

The time to extinction *t*_*ext*_ exhibits a similar trend. On average, the communities with more “complex” phenotypes go extinct faster when subjected to changing environmental conditions, and we observe this effect for small and for large values of *V*_*C*_ (Fig. 2F). Moreover, the extinction threshold becomes lower for increasing phenotypic complexity, which means that low-dimensional communities are able to withstand rates of environmental changes that would lead to extinction of the whole community in high-dimensional phenotype spaces (Appendix, Fig. S3B). A mechanistic explanation for the observed reduction of the extinction threshold is that in higher dimensions, the population size *N*_*r*_ of each species in the community is generally smaller. In higher-dimensional systems, each species has on average more competitors due to larger number of “nearest neighbours” with slightly different phenotypes. Smaller population sizes mean lower per species mutation rates, and hence slower adaptation to changing environments (Eq. 6).

To check the robustness of these results, we have repeated these simulations for fewer than 30 replicas for two other values of the width of the competition kernel, *σ*_*i*_ = 0.25 and 0.75. Even though the saturated level of diversity varies strongly with *σ_i_* (Doebeli and Ispolatov 2017), the trends shown and Figs. 2 and 3 remain qualitatively unchanged.

## Discussion

We have investigated the evolutionary dynamics in logistic competition models under the assumption of gradual environmental change, which was implemented by assuming that the maximum of the carrying capacity function moves at a constant speed in phenotype space. We have analyzed the effect of such environmental change on various statistical ecological and evolutionary properties of the adapting communities, depending on the rate of environmental change and on the dimension of phenotype space.

In our models, the crucial factor determining whether an evolving community will survive in the long run is the rate of environmental change. We found that there is always a threshold rate of environmental change above which communities cannot survive. However, when the rate of change is below the threshold value, many communities are able to adapt to environmental changes, and the evolving community can find a new quasi-stationary state. Such adaptation requires much less time compared to the time it takes a community to reach the pre-change level of diversity from a single ancestral species. To adapt, the phenotype of a species generally has to evolve in step with the movement of the centre of the carrying capacity, because otherwise the carrying capacity of that species will fall to very low values, making the environment inviable for that species. Such effects can be observed in various populations (Gienapp et al. 2008; Baym et al. 2016; Tseng et al. 2018). For example, numerous studies indicated a global tendency of various species to decrease in body size in warming environments (Guillemain et al. 2005; Y. Yom-Tov and J. Yom-Tov 2005; Y. Yom-Tov and S. Yom-Tov 2006; Y. Yom-Tov, Leader, et al. 2010; Gardner et al. 2011). Body size is an important physiological characteristic that strongly influences metabolism and thermoregulation. For example, decreased body-size due to changing temperature regimes was observed in beetles both in natural and laboratory populations (Tseng et al. 2018).

The rate of environmental change also affects the composition of newly formed quasi-stationary states. The higher the rate of environmental change is, the lower is the average number of coexisting species and the total population size of surviving communities. However, surviving species get an ecological advantage: because of the reduction of species competing in the community, the populations of each surviving species can become larger than in the case of stable environmental conditions. Thus, generally the fewer species are left in the surviving community, the larger their population size.

Interestingly, phenotypic complexity makes communities less resilient to changes in the environment. In high-dimensional phenotypic spaces surviving systems require more time to adapt, and hence are less able to resist changes, and the extinction threshold, below which no systems survive, appears at lower rates of environmental change. According to our previous studies of macroevolutionary processes (Doebeli and Ispolatov 2014, 2017; Ispolatov, Alekseeva, et al. 2019), “complex phenotypes” have smaller populations and lower rates of evolution, which makes adaptation challenging. In these models, evolution under constant environmental conditions results in gradual expansion of phenotypic space, and more complex, higher-dimensional phenotypes evolve only once diversity in lower dimensions has saturated (Ispolatov, Alekseeva, et al. 2019). Conversely, changing environments tend to reduce the number of phenotypic dimensions, since less complex species are more likely to survive. However, evolving communities that have become less diverse due to the environmental change may rediversify and reach previous saturation levels if the environment ceases to change. If that happens, the new phenotypic composition would gradually converge to the one existing before the onset of environmental change, while the genealogical history and composition of the re-diversified community may be very different from the composition of the community that existed before the environmental change was initiated.

Finally, it is worth noting that environmental changes have led in the past to more cataclysmic evolutionary changes, such as the evolution of entirely novel phenotypes. In terms of our models, such environmental changes would then likely generate new bouts of rapid diversification leading to saturated communities occupying higher dimensional phenotype spaces. In the future we plan to include such dramatic evolutionary events in our models.

## Acknowledgement

I.I. acknowledges support from DICYT project 041931Y. E.A. acknowledges support from SkolTech Academic Mobility Program.

## Data Accessibility

codes and averaged results of simulations are available at https://github.com/EvgeniiaAlekseeva/Climate.

## Appendix

## 1. Simulation procedure

**Figure S1.**
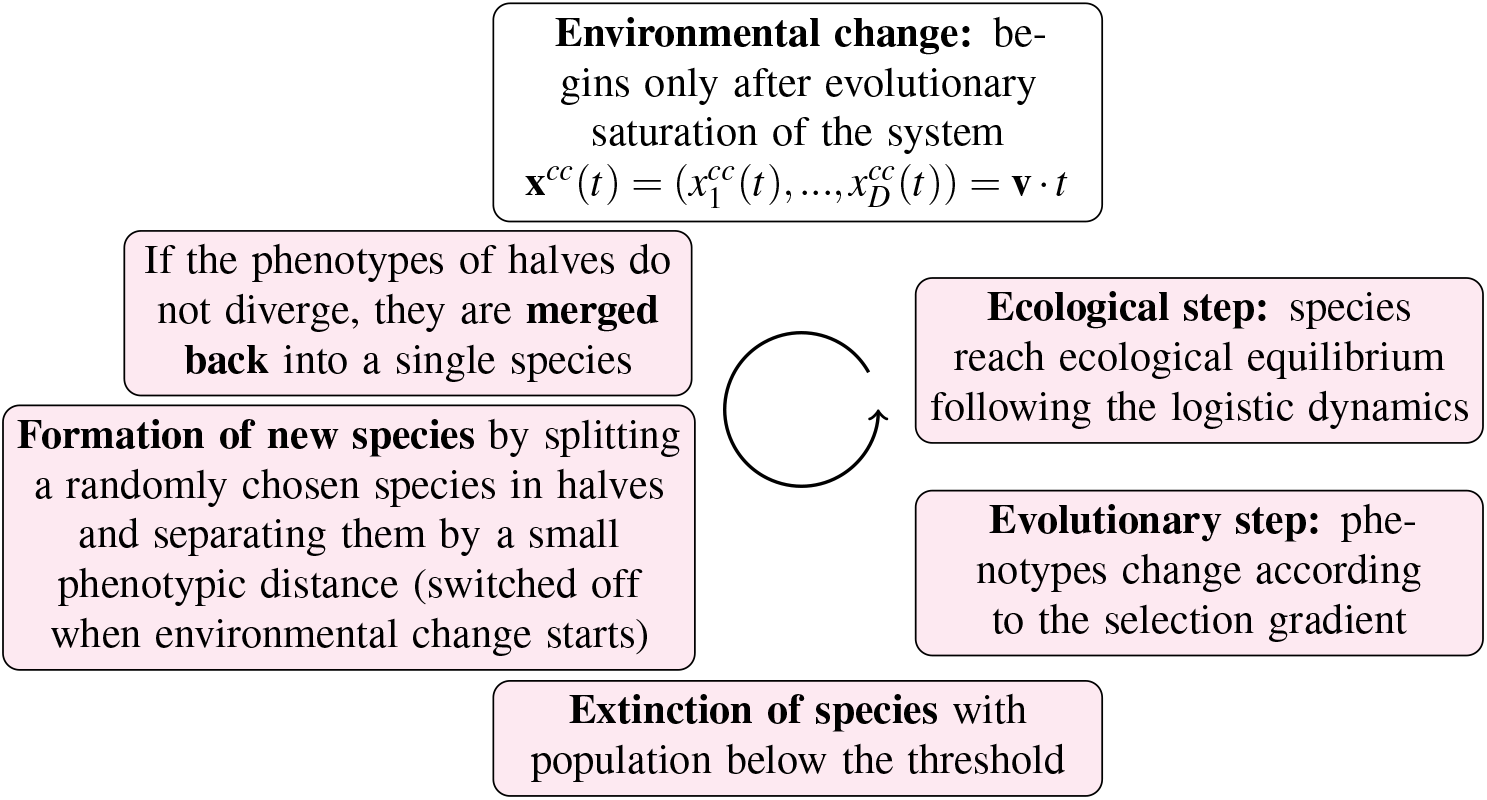
Successive steps that are iterated in the simulations. Each iterative cycle advances the evolutionary time by a small increment. The merging and splitting steps are performed once every 10 time units during the evolutionary saturation of the system. Environmental change (movement of CCC) starts only after the system converges to a steady state with saturated diversity and the formation of new species stops.

Simulations consist of many cycles of successive steps (Fig. S1), where each cycle increments the evolutionary time by a small amount. An iteration starts with the ecological dynamics, where all species reach their ecological equilibrium according to the logistic dynamics (Eq. 3). Evolutionary time stays constant during this step. If a species crosses the low population limit set equal to 10^−6^, it is assumed to be extinct and is dropped from the system. In the next step the phenotypes of all species evolve according to the adaptive dynamics specified in Eq. 6. Phenotypic changes in a single evolutionary time step Δ*t* ~ 10^−2^ are small enough to keep populations close to their ecological equilibrium (calculated in the previous step).

To model diversification, each 10 time units we split a randomly chosen species in halves separated by a very small phenotypic distance Δ*x* (normally Δ*x* = 10^−3^). If conditions are favorable for evolutionary branching, the distance between the halves grows as a result of phenotypic dynamics, and the two “halves” become two separate species. Otherwise, i.e., if the competitive interactions do not favor diversification and the halves do not move apart phenotypically, we merge them back without any consequences to system behavior. Both procedures of merging and splitting happen regularly every few iterations (in this order, so that split halves have time to diverge). In principle, if any two “not closely related” species come close in phenotypic distance at the time of merging, they would be merged as well, however, we have not observed such events in our simulations.

An evolving system is initiated as a single species with a phenotype that is randomly chosen from a Gaussian distribution with width 1 and mean 0. The evolutionary equilibration time *t*\* is defined as the time it takes the system to evolve from a single species to the saturated diversity. This time *t*\* quickly grows with the dimension of phenotypic space (Fig. S2). The procedure of evolving a system to saturated diversity is described in details in Doebeli and Ispolatov 2017.

In the principal part of our simulations, we analyzed how saturated systems adapt to environmental changes, moving the center of the carrying capacity with various speeds *V*_*C*_ and observing the changes in the systems over an evolutionary time period equal to *t*\* or until all species become extinct. Before the beginning of environmental changes, we merge all clusters with phenotypes within a much larger merging distance than used in the diversification procedure (usually set to 10^−1^) into distinct species, visible as circles in Fig. 1 and corresponding videos. This is done to ensure that the population density *N*_*r*_, which controls the adaptive dynamics evolutionary speed shown in Eq. (6), corresponds to the intergral population of the species, rather than to the “metapopulation” of each cluster. A “cloud” of such clusters, separated by phenotypic distance marginally larger than the merging distance Δ*x* = 10^−3^, is often created by our diversification procedure. The biological motivation behind this final merge is that each of the individual clusters can produce a mutant that could take over the whole cloud of those neighbouring clusters, i.e, the whole species. Using the usually smaller individual metapopulations as the factor *N*_*r*_ in Eq. (6), would have reduced the evolutionary speed and led to unrealistically more difficult adaptation and earlier extinction. Furthermore, starting from the onset of environmental change, we switch off the procedure of new species formation, as we do not expect the diversification of species under the pressure of environmental change.

The duration of simulation *t*\* was chosen to be the same as the time it takes a system to reach diversity saturation before the onset of environmental change. If a system survives the effects of CCC motion for the time *t*\*, it usually means that it reaches a new evolutionary steady state adapted to the constant environmental change.

**Figure S2.**
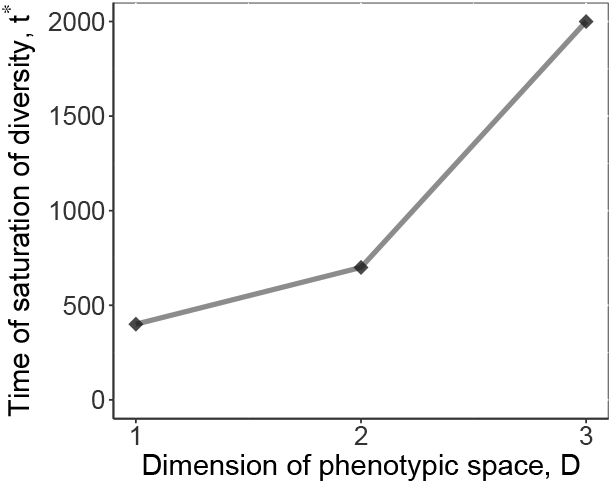
The evolutionary time *t*\* required for diversity saturation starting from the initial condition of one randomly located species. Around the time *t*\*, the number of species *m*_*tot*_ equilibrates.

The range of CCC speed *V*_*C*_ can be capped using the following simple analytical estimate for the maximum evolutionary speed that a single species could sustain. The case of a single surviving species is often the final outcome of adaptation to sufficiently rapid environmental changes. If we ignore the asymmetric part of the competition kernel represented by the random coefficients *b*_*ij*_ in Eq. (6), which could either reduce or increase the strength of competition in a generally unpredictable way, the remaining adaptive dynamics becomes quite simple. The evolutionary speed **u** has components

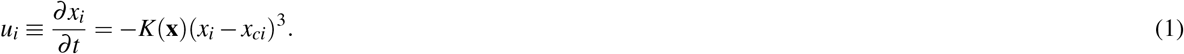

Here we have taken into account that the ecologically equilibrated single-species population is equal to the corresponding carrying capacity. Assuming for simplicity that the CCC moves along the first phenotypic coordinate, we look for the maximum of *u*_1_, differentiating (1) with respect to *x*_*i*_ −*x*_*ci*_. The maximum is achieved at 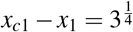, and the corresponding maximum speed of species motion in phenotypic space is *u*_*max*_ = (3/*e*)^3/4^ ≈ 1.08. This sets the upper limit on the sustainable velocity of CCC, which results from a combination between two trends: A faster motion of CCC makes the species to trail further behind in phenotype space, thus generating a larger selection gradient. However, the further a species trails behind the CCC, the lower is its population, which makes mutations more rare. The combination of these two trends defines the maximum velocity at which the single species can evolve, which, in other words, is the maximum velocity of CCC that a species can follow at a steady state. In reality, due to the action of the asymmetric terms in the competition kernel, and due to interspecies competition, some species go extinct below this maximum CCC velocity, while others persist even in slightly faster changing environments. The latter happens when a particular combination of randomly chosen *b*_*ij*_ coefficients results in a stronger selection gradient in the direction of the *V*_*C*_.

## 2. Visualized simulation run

Visualized simulations, presented in Fig.1 and available for download <u>here</u> have the following parameters and initial conditions:

(A) : Diversity saturation

Simulations starts from a single species with population *N* = [0.3] and phenotypic coordinates

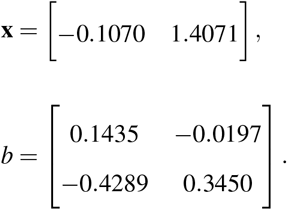

**(B,C,D)**: Adaptation to environmental changes

Simulations starts from the steady state of a simulation (**A**), consisting of *m* = 9 species with populations *N* = [0.4819 0.4923 0.4501 0.5187 0.5092 0.4704 0.47321 0.4432 0.4726] and phenotypic coordinates

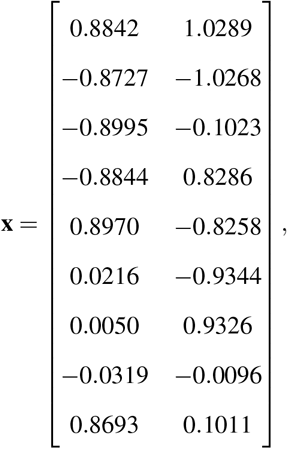

## 3. Threshold value of *V*_*C*_

**Figure S3.**
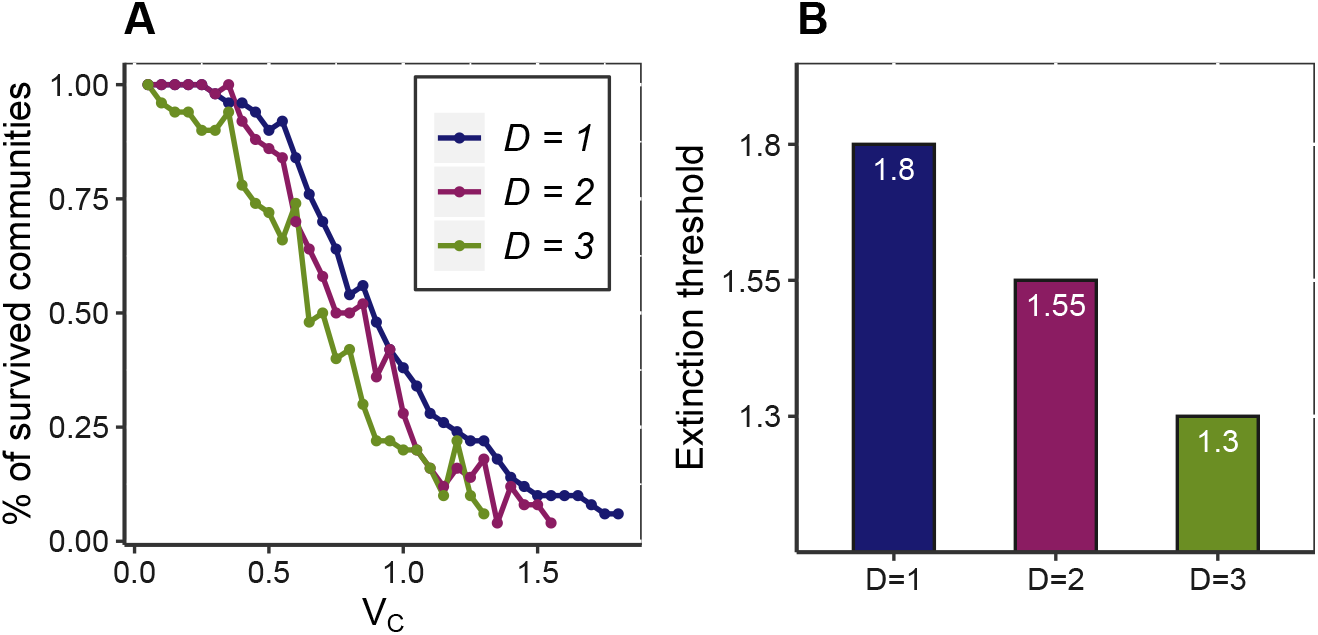
A: The fraction of systems with at least one surviving species from the sample of 30 saturated systems vs. the speed of CCC motion for three dimensions of phenotypic space; **B**: Extinction threshold 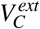 for three dimensionalities of phenotypic space; **C** The fraction of surviving species in the evolving community, ***ρ***_*surv*_; Surviving communities were able to adapt to the motion of CCC and reached a new quasi-stationary state.

